# (–)-Epicatechin alters reactive oxygen and nitrogen species production independent of mitochondrial respiration in human vascular endothelial cells

**DOI:** 10.1101/2021.10.25.465611

**Authors:** Daniel G. Sadler, Jonathan Barlow, Richard Draijer, Helen Jones, Dick H. J. Thijssen, Claire E. Stewart

## Abstract

**Introduction:** Vascular endothelial dysfunction is characterised by lowered nitric oxide (NO) bioavailability, which may be explained by increased production of reactive oxygen species (ROS), mitochondrial dysfunction and altered cell signalling. (-)-Epicatechin (EPI) has proven effective in the context of vascular endothelial dysfunction, but the underlying mechanisms associated with EPI’s effects remain unclear.

**Objective(s):** Our aim was to investigate whether EPI impacts reactive oxygen and nitrogen species (RONS) production and mitochondrial function of human vascular endothelial cells (HUVECs). We hypothesised that EPI would attenuate ROS production, increase NO bioavailability, and enhance indices of mitochondrial function.

**Methods:** HUVECs were treated with EPI (0-20 µM) for up to 48 h. Mitochondrial and cellular ROS were measured in the absence and presence of antimycin A (AA), an inhibitor of the mitochondrial electron transport protein complex III, favouring ROS production. Genes associated with mitochondrial remodelling and the antioxidant response were quantified by RT-qPCR. Mitochondrial bioenergetics were assessed by respirometry and signalling responses determined by western blotting.

**Results:** Mitochondrial superoxide production without AA was increased 32% and decreased 53% after 5 and 10 µM EPI treatment vs. CTRL (*P*<0.001). With AA, only 10 µM EPI increased mitochondrial superoxide production vs. CTRL (25%, *P*<0.001). NO bioavailability was increased by 45% with 10 µM EPI vs. CTRL (*P*=0.010). However, EPI did not impact mitochondrial respiration. NRF2 mRNA expression was increased 1.5- and 1.6-fold with 5 and 10 µM EPI over 48 h vs. CTRL (*P*=0.015 and *P*=0.001, respectively). Finally, EPI transiently enhanced ERK1/2 phosphorylation (2.9 and 3.2-fold over 15 min and 1 h vs. 0 h, respectively; *P*=0.035 and *P*=0.011).

**Conclusion(s):** EPI dose dependently alters RONS production of HUVECs but does not impact mitochondrial respiration. The induction of NRF2 mRNA expression with EPI might relate to enhanced ERK1/2 signalling, rather than RONS production. In humans, EPI may improve vascular endothelial dysfunction via alteration of RONS and activation of cell signalling.

## Introduction

Globally, cardiovascular disease (CVD) is the leading cause of morbidity and mortality (WHO, 2018). One major risk factor for CVD is vascular endothelial dysfunction, which is typified by impaired vasodilation and diminished blood flow (Heitzer et al., 2001; Suwaidi et al., 2000). Several factors contribute to vascular endothelial dysfunction, including reduced nitric oxide (NO) bioavailability and elevated oxidative stress (Ungvari et al., 2018). A decline in NO bioavailability has been attributed to lower endothelial nitric oxide synthase (eNOS) content and activity, in part due to lower phosphorylation of eNOS at Ser1177 - at least in the aged vascular endothelium (Gliemann et al., 2018; Musicki et al., 2005). Elevated oxidative stress might be explained increased production of ROS (Cernadas et al., 1998; Chou et al., 1998; Donato et al., 2007, 2009), that are cytosolic and mitochondrial in origin (Csiszar et al., 2002, 2007; Donato et al., 2007; Hamilton et al., 2001; Jablonski et al., 2007; Sun et al., 2004; Van Der Loo et al., 2000).

Vascular endothelial health is also impacted by mitochondrial function. Indeed, ageing is associated with reduced mitochondrial content in endothelial cells of conduit arteries, feed arteries and capillaries (Burns et al., 1979; S. H. Park et al., 2018; S.-Y. Park et al., 2018; Ungvari et al., 2008), which could be due to blunted transcriptional responses (S.-Y. Park et al., 2018; Ungvari et al., 2008). This potential reduction in mitochondrial biogenesis with ageing may result from diminished NO bioavailability (Gouill et al., 2007; Miller et al., 2013) and/or lowered AMP-activated protein kinase (AMPK) signalling (Lesniewski et al., 2012), both of which are known to activate peroxisome proliferator-activated receptor gamma coactivator 1-alpha (PGC-1α). A link between mitochondria and vascular endothelial dysfunction has also been made from observations that human skeletal muscle feed arteries exhibit impaired respiratory capacity and lower coupling efficiency in middle-(55 years) and older-age (70 years) compared to young adults (S. H. Park et al., 2018; S.-Y. Park et al., 2020). Furthermore, mitochondrial-targeted antioxidants like Mitoquinone (MitoQ) restore vascular endothelial dysfunction in aged mice and patients with peripheral artery disease, which likely results from reductions in levels of mitochondrial superoxide (Gioscia-Ryan et al., 2014; S.-Y. Park et al., 2020). Together, evidence points towards mitochondria as promising targets for interventions aimed at combatting vascular endothelial dysfunction.

(-)-Epicatechin (EPI) belongs to a subclass of flavonoids known as the flavanols. Not only is EPI highly bioavailable – reaching up to 10 µM in circulation in humans (Hollands et al., 2013) - but EPI is also associated with several health benefits (Arts et al., 2001; Hertog et al., 1993), including improved vascular endothelial function (Galleano et al., 2013; Karim et al., 2000; Schroeter et al., 2006) and increased NO bioavailability (Schroeter et al., 2006). Additional mechanisms thought to underly the therapeutic effects of EPI include: increased eNOS phosphorylation (Ramirez-Sanchez et al., 2011, 2012, 2018), enhanced content or activity of enzymatic antioxidant proteins (Moreno-Ulloa, Nogueira, et al., 2015; Ramirez-Sanchez et al., 2014) and augmented mitochondrial biogenesis (Lee et al., 2015; Ramirez-Sanchez et al., 2018; Taub et al., 2012, 2016). Whilst the potential of EPI to enhance markers of mitochondrial biogenesis and antioxidant capacity seems promising, it is unclear whether this effect translates to enhanced respiratory function or altered ROS production of vascular endothelial cells. In-fact, one recent study reported no impact of EPI on mitochondrial respiration (Keller et al., 2020), but the authors demonstrated that EPI may lower mitochondrial ROS production of HUVECs in the presence of high glucose. The challenge remains to resolve whether EPI modulates ROS production and mitochondrial function of vascular endothelial cells.

To this end, our aim was to investigate whether EPI modulates reactive oxygen and nitrogen species (RONS) production and mitochondrial function of human vascular endothelial cells. We hypothesised that EPI would attenuate ROS production, augment NO bioavailability, and enhance indices of mitochondrial function.

## Materials and Methods

### Cell culture and treatment

Human umbilical endothelial vein endothelial cells (HUVECs; Thermo Fisher Scientific, Waltham, MA, USA) at passages 3-7 were used in this study. HUVECs were not passaged more than 7 times because of changes in cell phenotype that can occur with multiple population doublings that ultimately lead to senescence (Chang et al., 2005; Cheung, 2007; Grillari et al., 2000). Following the plating of cells onto pre-gelatinised well-plates (0.2% gelatin) in complete endothelial cell growth medium (EGM; Cell Applications Inc, San Diego, CA, USA), ∼80% confluent HUVECs were washed twice with Dulbecco’s phosphate-buffered saline (D-PBS) and switched to pre-warmed (37°C) EGM in the absence (vehicle [H_2_O], ‘CTRL’) or presence of EPI (0.5-20 µM) over 24 h and 48 h. Human umbilical endothelial vein endothelial cells (HUVECs; Thermo Fisher Scientific, Waltham, MA, USA) at passages 3-7 were used in this study. HUVECs were not passaged more than 8 times because of changes in cell phenotype that can occur with multiple population doublings that ultimately lead to senescence (Chang et al., 2005; Cheung, 2007; Grillari et al., 2000). Following the plating of cells onto pre-gelatinised well-plates (0.2% gelatin) in complete endothelial cell growth medium (EGM; Cell Applications Inc, San Diego, CA, USA), ∼80% confluent HUVECs were washed twice with Dulbecco’s phosphate-buffered saline (D-PBS) and switched to pre-warmed (37°C) EGM in the absence (vehicle [H_2_O], ‘CTRL’) or presence of EPI (0-20 µM) over 24 h and 48 h.

### Cell viability

The fluorescent CyQUANT^®^ Proliferation Assay kit was used to determine cell viability. HUVECs were grown to 60-70% confluency in EGM in gelatinised 96-well plates. Cells were subsequently dosed for 24 h in EGM +/-EPI at 0-20 µM. After 24 h, wells were aspirated, washed twice with D-PBS, and then frozen immediately at -80°C. On the day of the assay, plates were thawed at room temperature and CyQUANT^®^ GR dye/cell-lysis buffer was added to each well according to manufacturer instructions. Plates were gently mixed on an orbital shaker (80 rpm) for 5 minutes protected from light. Sample fluorescence was measured using a CLARIOStar plate reader (BMG Labtech, Ortenberg, Germany) with 485/520 Ex/Em.

### Mitochondrial ROS production

Mitochondrial superoxide was detected in HUVECs using MitoSOX (Thermo Fisher Scientific, Waltham, USA), a hydroethidine probe which is targeted to mitochondria by a conjugated triphenyl-phosphonium moiety (Zielonka et al., 2017). HUVECs were seeded at 3 × 10^4^ cells/mL in gelatinised 12-well microplates and at ∼80% confluence dosed +/-EPI (0-10 µM) for 24 h. Next, cells were washed in Krebs-Ringer buffer (KRH; 135 mM NaCl, 3.6 mM KCl, 10 mM HEPES (pH 7.4), 0.5 mM MgCl_2_, 1.5 mM CaCl_2_, 0.5 mM NaH_2_PO_4_, 2 Mm glutamine and 5 mM D(+)-Glucose) prior to incubation at 37°C for 30 minutes, with or without 15 µM antimycin A (AA) to stimulate mitochondrial superoxide production. Next, AA-containing KRH was removed and MitoSOX was loaded into cells in fresh pre-warmed KRH to a final concentration of 2.5 µM. Plates were immediately transferred to a plate reader (ClarioStar, BMG Labtech) and fluorescence was monitored continuously at 30-sec intervals over 30 min at excitation/emission of 510/580 nm. Rates of mitochondrial superoxide production were determined from the slope of the resultant progress curve over the 30-minute recording. Finally, plates were immediately fixed in 1% (v/v) acetic acid in methanol for the determination of cell density by the Sulforhodamine B (SRB) assay, which was used to normalise obtained fluorescence values. The plate reader’s focal height and gain were optimised and fixed between different experiments.

### Cellular ROS

Cellular ROS were detected using the CellROX^®^ Deep Red reagent by spectrophotometry. HUVECs were seeded at 3 × 10^4^ cells/mL into gelatinised 12-well microplates and at ∼80% confluence dosed +/-EPI (0-10 µM) for 24 h. After treatment, HUVECs were washed in KRH with or without 15 µM AA and incubated at 37°C for 30 minutes, prior to KRH removal and CellROX loading using fresh, pre-warmed KRH buffer, to a final concentration of 2.5 µM. Following 30 minutes CellROX incubation protected from light, cells were washed 2 × with D-PBS and immediately transferred to a plate reader (ClarioStar, BMG Labtech), where fluorescent CellROX oxidation products were excited at 640 nm and light emission detected at 665 nm. The plate reader’s focal height and gain were optimised and fixed between experiments. Upon completion of the reading, plates were immediately fixed for the determination of cell density by as discussed above.

### Nitric oxide bioavailability

To assess intracellular NO bioavailability, HUVECs were plated and cultured as previously described. At ∼80% confluency, cells were treated with 0, 5 or 10 µM EPI for 24 h. After treatment, HUVECs were washed 2 × with D-PBS and loaded with DAF-FM™ diacetate (4-amino-5-methylamino-2′,7′-difluorofluorescein diacetate; Molecular Probes, Invitrogen) to a final concentration of 1 µM in KRH buffer and incubated at 37°C for 45 minutes protected from light. Following dye loading, cells were washed 2 × with D-PBS and immediately trypsinised prior to pelleting and resuspension in D-PBS. Sample fluorescence was subsequently detected at 495/515 Ex/Em by flow cytometry (BD Accuri C6, BD Biosciences, Wokingham, UK). Data were recorded from 5,000 events and median values reported.

### RT-qPCR – Gene expression quantification

HUVECs were grown to ∼70% confluency in EGM in gelatinised 12-well plates and lysed in 125 µL TRIzol. Total RNA was then extracted using the phenol-chloroform method. RNA concentrations were determined by spectrophotometry (NanoDrop™ 2000, Thermo Fisher Scientific, Waltham, USA). Specific primers used in each PCR are outlined in supplementary Table 1. After preparation, reaction tubes were transferred to a Rotor-Gene Q PCR thermal cycler for product amplification using a one-step protocol (QuantiFast SYBR^®^ Green RT-PCR Kit, Qiagen, UK). The amplification protocol was as follows: reverse transcription (10 minutes at 50°C), transcriptase inactivation and initial denaturation (95°C for 5 min) followed by 40 × amplification cycles consisting of: 95°C for 10 s (denaturation) and 60°C for 30 s (annealing and extension); followed by melt curve detection. Critical threshold (C_T_) values were derived from setting a threshold of 0.09 for all genes. To quantify gene expression, C_T_ values were used to quantify relative gene expression using the comparative Delta Delta C_T_ (2^-ΔΔCT^) equation (Livak & Schmittgen, 2001), whereby the expression of the gene of interest was determined relative to the internal reference gene (RPL13a) in the treated sample, compared with the untreated zero-hour control.

**Table 1.** List of antibodies and dilutions used.

| Antibody | Primary Ab Dilution | Secondary Ab Dilution |
| --- | --- | --- |
| AMPKα | 1:1000 | 1:5000 |
| pThr172-AMPK | 1:1000 | 1:10,000 |
| p44/42 MAPK | 1:2000 | 1:10,000 |
| pThr202/Tyr204-p44/42<br>MAPK | 1:2000 | 1:10,000 |
| eNOS | 1:500 | 1:5000 |
| pSer1177-eNOS | 1:500 | 1:5000 |

All antibodies were purchased from Cell Signaling Technology.

### Mitochondrial respiration

HUVECs (passages 4-6) were seeded in XFe 24 well plates (Agilent, Santa Clara, CA, USA) at 30,000 cells per well in 200 µL EGM for 48 h. After 48 h, HUVECs were washed twice with D-PBS and replaced with fresh EGM containing 0, 5 and 10 µM EPI for 24 h. On the day of the assay, HUVECs were washed with pre-warmed XF Dulbecco’s Modified Eagle Medium pH 7.4 (DMEM; Agilent, Santa Clara, CA, USA), supplemented with 5.5 mM glucose, 1 mM sodium pyruvate and 2 mM L-Glutamine, and brought to a final well volume of 500 µL. The cells were incubated in this medium for 45 minutes at 37°C in a non-CO_2_ incubator and then transferred to a Seahorse XFe24 extracellular flux analyser (Agilent, Santa Clara, CA, USA) maintained at 37°C. After an initial 15-minute calibration, oxygen consumption rate (OCR) was measured by a 3-4 loop cycle consisting of a 1-min mix, 2-min wait and 3-min measure to record cellular respiration. After measuring basal respiration, 2 mM oligomycin was added to selectively inhibit the mitochondrial ATP synthase. Subsequently, 3 µM BAM15 was added to uncouple OCR to determine maximal respiration, and finally a mixture of 2 µM rotenone and 2 µM AA was added to inhibit complex I and III of the electron transfer system, respectively, to determine non-mitochondrial respiration. Rates of oxygen consumption were corrected for non-mitochondrial respiration and expressed relative to the cell number of the appropriate well, determined by the CyQUANT^®^ assay. The raw values of extracellular acidification rate (ECAR) and OCR were divided into component rates to calculate the relative contribution of glycolytic (ATP_glyc_) and oxidative ATP-producing reactions (ATP_ox_) to total ATP production, as previously described (Mookerjee & Brand, 2015). Three independent experiments were performed that contained at least two technical replicates.

### SDS-PAGE and immunoblotting

Total protein and phosphoprotein levels were detected by western blot. Following treatment, HUVECs were lysed and scraped in ice-cold 1x precipitation assay buffer containing: 25 mM Tris-HCl pH 7.6, 150 mM NaCl, 1% NP-40, 1% sodium deoxycholate and 0.1% SDS, supplemented with 1x Protease Inhibitor Cocktail Set V (Merck Life Science, UK). Cell lysates were centrifuged for 15 minutes at 18,000 × g (4°C) and the supernatant was stored at -80°C before analyses for total protein by the Pierce BCA™ assay. Samples were subsequently resuspended in 4x Laemmli buffer (Bio-Rad laboratories, Hertfordshire, UK) containing reducing agent (1x working concentration: 31.5 mM Tris-HCl [pH 6.8], 10% glycerol, 1% SDS, 0.005% Bromophenol Blue and 355 mM 2-mercaptoethanol) and were loaded and electrophoresed on 10% SDS-stain-free polyacrylamide gels (supplementary Figure 1). Semi-dry transfer of proteins to a nitrocellulose membrane was performed using the Trans-Blot^®^ Turbo™ Transfer System. Following blocking for 1-hour in Tris-buffered saline Tween-20 (TBS-T) containing 5% non-fat dried milk (NFDM), membranes were incubated overnight with rabbit anti-phosphorylated or total antibodies: AMPKα, pThr172-AMPK, p44/42 MAPK, pThr202/Tyr204-p44/42 MAPK, eNOS and pSer1177-eNOS, at a dilution of 1:500-1:4000 in 5% bovine serum albumin (BSA) made up in TBS-T (see Table 1). After overnight incubation, the membranes were washed 3 times in TBS-T for 5 minutes and incubated for 1 hour in HRP-conjugated anti-rabbit antibodies (Cell Signaling Technology, London, UK) at dilution of 1:5000-1:10,000, following optimisation. Proteins were visualised by enhanced chemiluminescence (Thermo Fisher Scientific inc, Waltham, USA) and quantified by densitometry (ChemiDoc™ MP imaging system, Bio-Rad Laboratories, Inc. CA, USA). Analysed western blot images are presented in supplementary figure 2.

### Statistical analysis

One-way ANOVAs were employed to detect effects of EPI treatment. Two-way ANOVAs were employed to detect main effects (e.g., dose, time or antimycin A) and potential significant interactions between two main independent factors. Multiple comparisons between experimental conditions were adjusted for multiple tests, using Dunnett’s or Sidak’s where appropriate. All data are presented as mean±SEM and significance accepted when *P*<0.05.

## Results

### EPI does not cause vascular endothelial cell toxicity

After 24 h EPI treatment (0.5-20 µM dose responses), there was a significant main effect of dose on cell proliferation (*P*=0.018; Figure 1). However, multiple comparisons revealed no significant difference between doses of EPI versus CTRL. Given that EPI did not cause cell toxicity, and prior knowledge of physiologic EPI concentrations *in vivo* (up to 10 µM), subsequent experiments were performed with doses of 5 and 10 µM.

**Figure 1.**
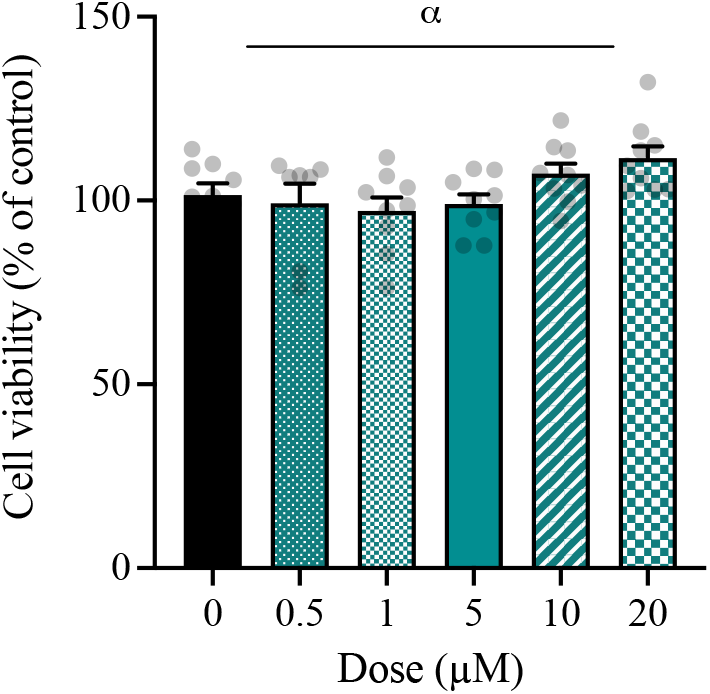
EPI does not cause **v**ascular endothelial cell toxicity. HUVECs were treated with 0-20 µM EPI for 24 h. Data are means±SEM, representative of 3 independent repeats with 3 replicates of each condition. Statistical significance was tested for by one-way ANOVA and Dunnett’s test for multiple comparisons. ^α^ Significant main effect of dose (*P*<0.05).

### EPI dose-dependently modulates mitochondrial RONS production

Next, we assessed whether EPI, in the absence or presence of the complex III inhibitor, AA, impacted mitochondrial superoxide emission. There was a significant main effect of dose and AA on rates of MitoSOX oxidation (*P*<0.001), and a significant dose × AA interaction (*P*<0.001; Figure 2A). Post-hoc comparisons revealed that, in the absence of AA (-AA), 5 µM EPI significantly increased and 10 µM EPI decreased rates of MitoSOX oxidation compared to CTRL, respectively (CTRL: 8.1×10^−5^±0.2×10^−5^; 5 µM EPI: 10.7×10^−5^±0.2×10^−5^; 10 µM EPI: 3.8×10^−5^±0.2×10^−5^ RFU/sec^-1^/cell^-1^; *P*<0.001). The associated raw traces of MitoSOX oxidation are displayed in Figure 2B. In the presence of AA (+AA), 5 µM EPI did not affect MitoSOX oxidation versus CTRL (5 µM EPI: 35.4×10^−5^±0.4×10^−5^ vs. CTRL: 35.4×10^−5^±0.5×10^−5^ RFU/sec^-1^/cell^-1^; Figure 2A). Whereas 10 µM EPI significantly increased rates of MitoSOX oxidation compared to CTRL (10 µM EPI: 44.4×10^−5^±0.6×10^−5^ vs. CTRL: 35.4×10^−5^±0.5×10^−5^ RFU/sec^-1^/cell^-1^; *P*<0.001). In contrast to mitochondrial ROS, cellular ROS production (not mitochondrial-specific) was not altered by EPI (Figure 2C).

**Figure 2.**
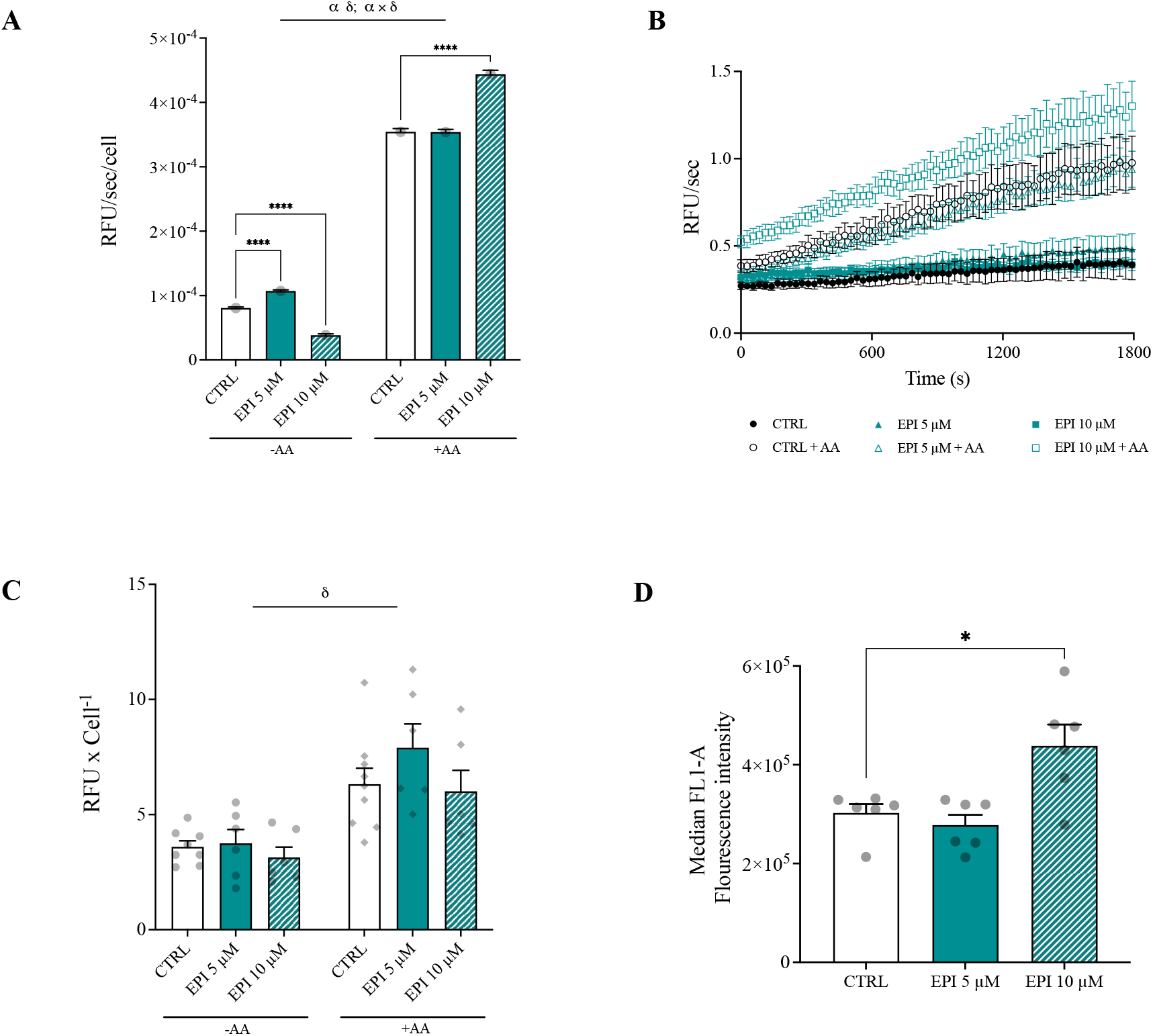
EPI dose-dependently impacts RONS production in vascular endothelial cells. A) MitoSOX oxidation rates in HUVECs after 24 h EPI treatment and 30 minutes incubation with or without antimycin A. B) Mean unnormalized MitoSOX oxidation rates measured in 30 second intervals over 30 minutes. C) CellROX oxidation in HUVECs after 24 h EPI treatment. D) DAF-FM oxidation in HUVECs after 24 h EPI treatment measured by flow cytometry. Data are means±SEM of three independent repeats with two replicates per treatment. Statistical significance was tested for by a two-way ANOVA, with dose and antimycin A as factors: ^α^ Significant main effect of dose; ^δ^ Significant main effect of AA (*P*<0.05). \**P*<0.05 and **** *P*<0.0001.

After revealing that rates of mitochondrial ROS production were dose-dependently altered by EPI, we assessed whether EPI modified intracellular NO levels. There was a significant main effect of EPI dose on NO levels (*P*=0.003; Figure 2D). Whilst 5 µM EPI did not impact NO levels, 10 µM EPI significantly increased NO levels compared to CTRL conditions (10 µM EPI: 4.38×10^5^±0.43×10^5^ vs. CTRL: 3.02×10^5^±0.18×10^5^ AU; *P*=0.010).

### EPI dose-dependently impacts the expression of genes associated with energy metabolism in vascular endothelial cells

Next, experiments were performed to resolve whether EPI alters the expression of genes linked with mitochondrial function and the antioxidant response. Firstly, the expression of genes associated with mitochondrial function and remodelling were quantified. In the presence of EPI, there was a significant main effect of dose (*P*=0.018) and time (*P*=0.002) on dynamin-related protein 1 (DRP1) expression (Figure 3A), but no significant dose × time interaction. At 48 h, 10 µM EPI increased DRP1 expression 2.2-fold compared to CTRL (*P*=0.010). There was a significant main effect of dose on mitofusin-2 (MFN2) mRNA expression in cells treated with EPI (*P*=0.035; Figure 3B). With 10 µM EPI, MFN2 expression increased 1.6-fold versus CTRL (*P*=0.024). Parkin, PGC-1α, and transcription Factor A (TFAM) expression were not changed by EPI treatment (supplementary Figure 3).

**Figure 3.**
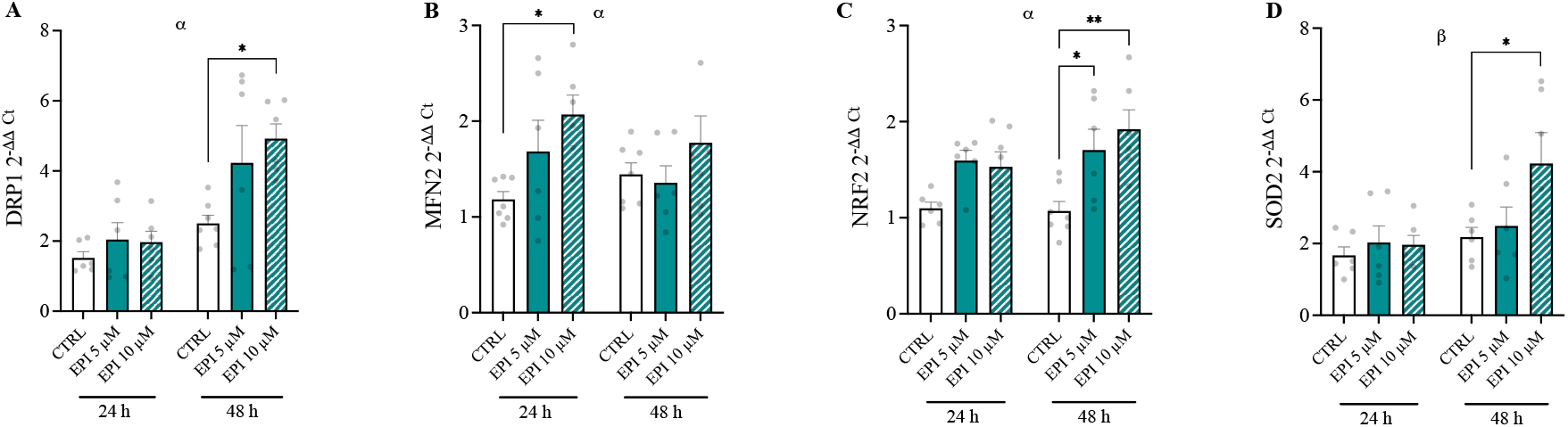
Gene expression responses following acute EPI treatment. HUVECs were treated with 0, 5 and 10 µM EPI over 48 h and lysed for analysis of gene expression. A) DRP1, B) MFN2, C) NRF2 and D) SOD2. Data are means±SEM from 3 independent experiments. Statistical significance was determined by a two-way ANOVA, with dose and time as factors. Multiple comparisons were performed by Dunnett’s test to determine differences in gene expression between conditions. ^α^ Main effect of dose; ^β^ main effect of time (*P*<0.05); \**P*<0.05, \*\**P*<0.01.

Genes associated with the antioxidant response were also quantified. There was no effect of dose or time on catalase or eNOS expression. There was a significant main effect of dose and time on NADPH oxidase 4 (NOX4) expression in EPI treated cells, respectively (*P*=0.015 and *P*=0.006). A significant main effect of dose was found on nuclear factor-erythroid factor 2-related factor 2 (NRF2) expression in the presence of EPI (*P*<0.001; Figure 3C). NRF2 mRNA abundance was increased 1.5-fold and 1.6-fold with 5 and 10 µM EPI over 48 h when compared to CTRL (*P*=0.015 and *P*=0.001, respectively). There was a significant effect of time on superoxide dismutase 2 (SOD2) expression in EPI treated cells only (*P*=0.024). Multiple comparisons revealed that SOD2 expression was increased 2.1-fold in the presence of 10 µM EPI versus CTRL conditions (*P*=0.040; Figure 3D).

### EPI does not alter mitochondrial bioenergetics of vascular endothelial cells

Having described that EPI influences RONS production and alters the expression of genes linked with mitochondrial function, we tested whether EPI impacted vascular endothelial cell bioenergetics. There was no significant main effect of EPI on rates of basal respiration, maximal respiration, ADP phosphorylation, proton leak, spare respiratory capacity (%) or coupling efficiency, regardless of dose (Figure 4A & 4B & 4D). There was no effect of EPI on the relative contribution of ATP_glyc_ or ATP_ox_ to total ATP production (Figure 4C).

**Figure 4.**
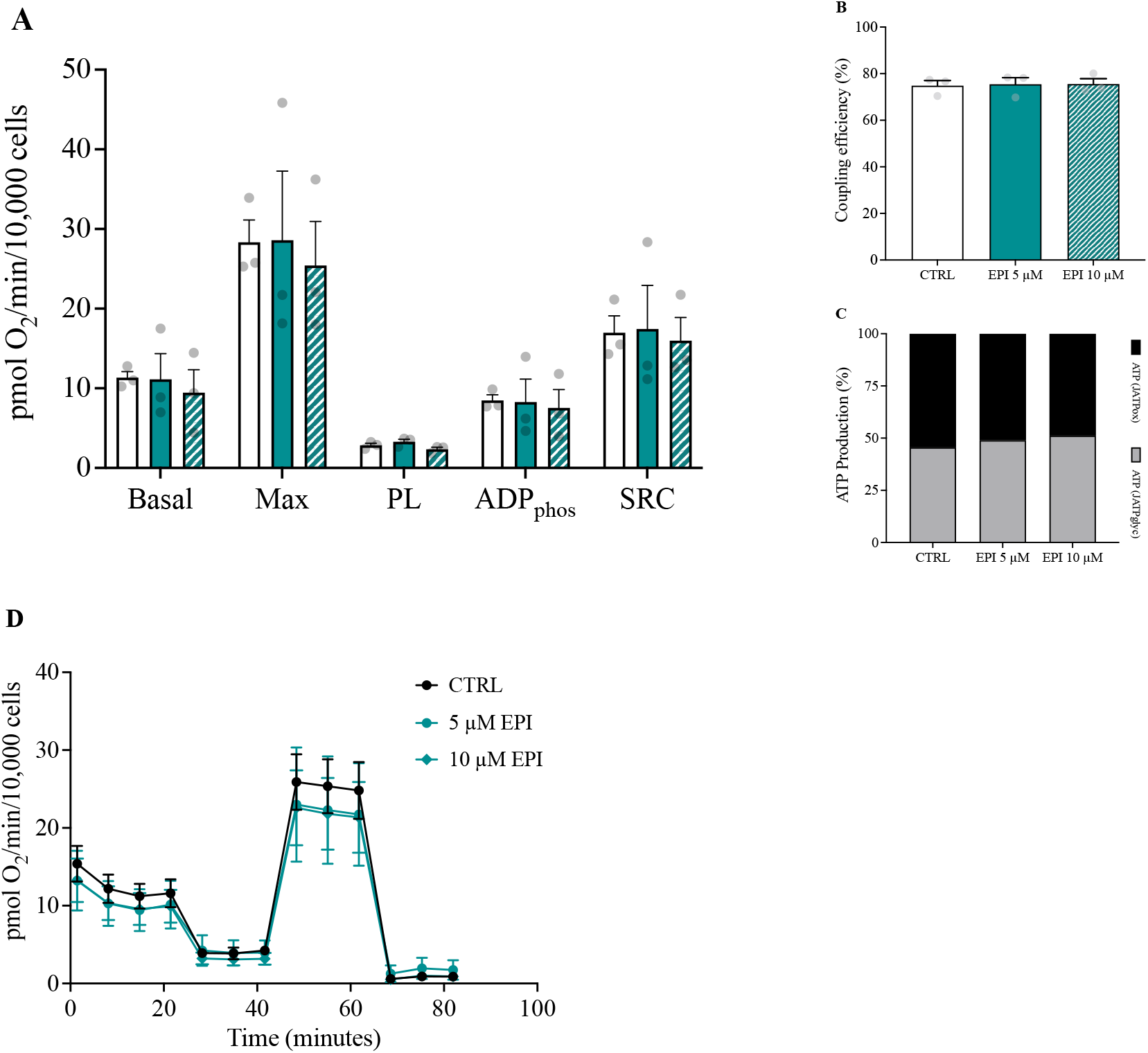
EPI does not directly affect mitochondrial bioenergetics. A) Mitochondrial bioenergetics of HUVECs following 24 h EPI treatment (0, 5 and 10 µM). B) Coupling efficiency of oxidative phosphorylation. C) Relative contribution of glycolytic (grey) and oxidative (black) ATP production to total ATP production rates. D) Representative trace of oxygen consumption rates during the mitochondrial stress test. Data from 3 independent experiments are normalised to cell number (1×10^3^) and presented as mean±SEM. Max, maximal respiratory capacity; PL, proton leak; ADPphos, ADP phosphorylation; SRC, spare respiratory capacity.

### EPI rapidly and transiently promotes ERK1/2 phosphorylation, independent of AMPK

To further probe how EPI might alter vascular endothelial transcription, we assessed cell signalling responses. There was a significant main effect of treatment and time on AMPKα phosphorylation, and a significant interaction was observed (*P*<0.001; Figure 5A). Multiple comparisons revealed a significant increase in phosphorylation of AMPKα at Thr172 at 1 h versus 0 h (1 h: 2.24±0.22 vs. 0 h: 1.16±0.19 AU) under CTRL conditions (*P*=0.006), whereas there was no significant change at 1 h versus 0 h with EPI (1 h: 0.47±0.18 vs. 0 h: 1.16±0.19 AU; *P*=0.157). However, there was a significant reduction in AMPKα phosphorylation in the presence of EPI vs. CTRL at 15 min (EPI: 0.35±0.09 vs. CTRL: 1.86±0.29 AU; *P*<0.001) and 1 h (EPI: 0.47±0.18 vs. CTRL: 2.24±0.22 AU; *P*<0.001). From 3 hours, AMPKα phosphorylation was suppressed under both CTRL (3 h: 0.54±0.08 and 24 h: 0.39±0.12 AU vs. 0 h: 1.16±0.19 AU; *P*=0.046 and *P*=0.026, respectively) and EPI conditions compared to 0 h CTRL (3 h: 0.37±0.08 and 24 h: 0.46±0.08 AU vs. 0 h: 1.16±0.19 AU; *P*=0.021 and *P*=0.049, respectively).

**Figure 5.**
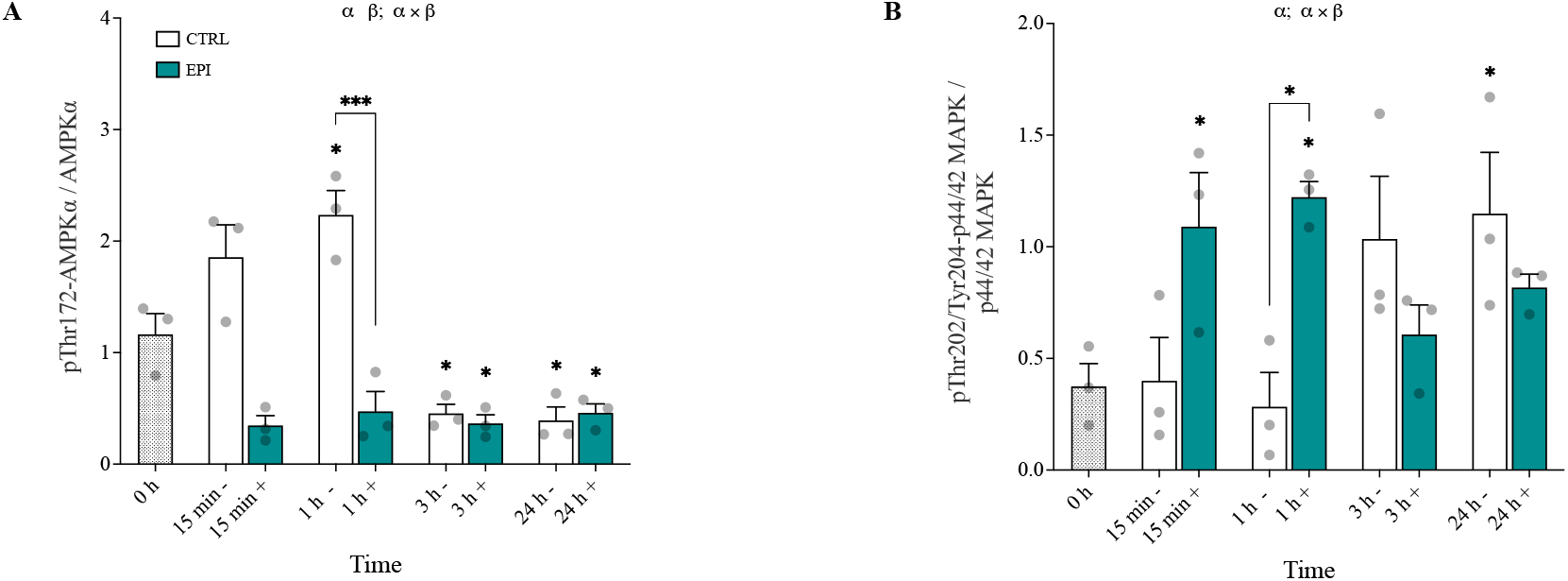
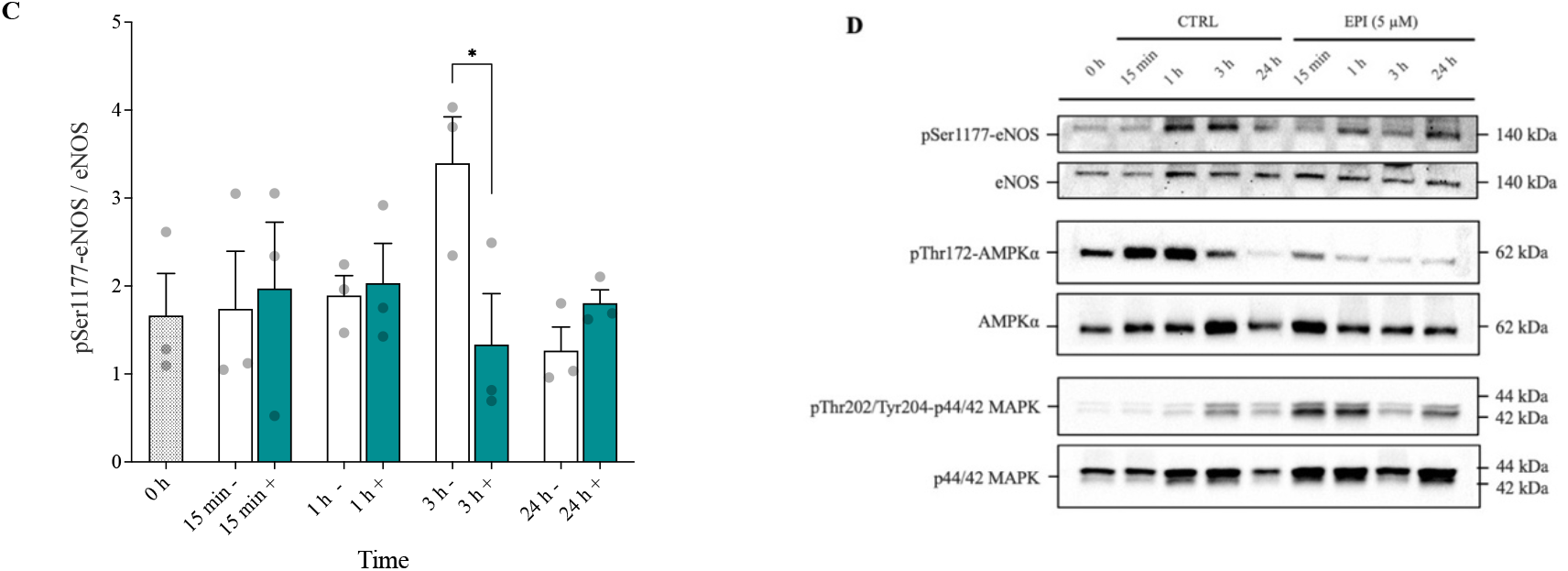
EPI rapidly and transiently activates ERK1/2 signalling whilst blunting AMPK phosphorylation. A) AMPKα phosphorylation at Thr172 in HUVECs in the absence (-; clear bars) or presence (+; green bars) of EPI. B) ERK1/2 phosphorylation at Thr202/Tyr204. C) eNOS phosphorylation at Ser1177. D) Representative images of n=3 independent experiments. Cell lysates were analysed by SDS-PAGE and western blotting with indicated antibodies. Data are expressed as means±SEM; \**P*<0.05 and \*\*\**P*<0.001. ^α^ Significant main effect of treatment; ^β^ significant main effect of time (*P*<0.05).

Whilst ERK1/2 is not involved in the AMPK/eNOS pathway, ERK1/2 signalling may mediate the effects of EPI upon vascular endothelial cell adaptation. There was no main effect of treatment (*P*=0.141), but a significant main effect of time on ERK1/2 phosphorylation (*P*=0.039; Figure 5B). There was also a significant treatment × time interaction (*P*=0.003). Under CTRL conditions, despite a 2.7-fold increase in ERK1/2 phosphorylation at 3 h, ERK1/2 phosphorylation did not reach significance vs. 0 h until 24 hours post treatment (0 h: 0.38±0.10 vs. 24 h: 1.15±0.28 AU; *P*=0.022). By contrast, EPI significantly increased ERK1/2 phosphorylation at 15 minutes vs. 0 h (15 min: 1.09±0.24 vs. 0 h: 0.38±0.10 AU; *P*=0.035), which was retained at 1 hour (1 h: 1.22±0.07 vs. 0 h: 0.38±0.10 AU; *P*=0.011), before returning to baseline levels, suggesting a change in the temporal pattern of ERK1/2 activation because of EPI treatment. Indeed at 1-hour, EPI treatment resulted in a 4.3-fold increase in ERK1/2 phosphorylation vs. the 1-hour untreated CTRL (*P*=0.007).

To help establish whether the increase in NO brought about by EPI was associated with eNOS signalling, phosphorylation of eNOS at Ser1177 was assessed. There was no main effect of treatment or time on eNOS phosphorylation (Figure 5C), and no treatment × time interaction (*P*=0.100). At 3 h, eNOS phosphorylation was ∼60% higher under CTRL versus EPI conditions (CTRL: 0.38±0.10 vs EPI: 1.15±0.28 AU, *P*=0.038).

## Discussion

Resolving EPI’s mode of action using vascular endothelial cells as a model will help to establish its potential efficacy in mitigating vascular endothelial dysfunction. We tested the hypothesis that EPI would attenuate ROS production, augment NO bioavailability and enhance mitochondrial function of human vascular endothelial cells in culture. We demonstrated that physiologic EPI concentrations, dose-dependently, modulated RONS emission but did not directly impact mitochondrial respiration. Moreover, the influence of EPI on RONS emission was associated with the induction of increased NRF2 mRNA, which appears downstream of rapid and transient activation of ERK1/2 signalling. Taken together our findings expand our knowledge of EPI’s mechanisms of action *in vitro* and support further research on EPI as a potential instigator of cell signalling and NRF2 activation *in vivo*.

### EPI dose-dependently modulates RONS production

We have demonstrated that EPI dose-dependently altered mitochondrial ROS production in the absence of additional cell stress. Increased rates of mitochondrial ROS production, reported in the presence of 5 µM EPI supports previous observations that 10 days EPI supplementation increased superoxide production in mitochondria isolated from mouse heart tissue (Panneerselvam et al., 2013). However, EPI has been demonstrated to increase the abundance and/or activity of key antioxidant proteins like SOD2 and catalase (Bettaieb et al., 2014; Calabró et al., 2016), and even to blunt hydrogen peroxide production from isolated brain and heart mitochondria (Lagoa et al., 2011). In a similar way, we found that higher (10 µM) EPI concentrations attenuated the rate of mitochondrial superoxide production, which may have been facilitated by increased SOD2 mRNA expression. Clearly and importantly, the divergent effects of EPI on mitochondrial ROS production suggest EPI’s biological effects are highly dose dependent. Future studies should investigate whether EPI alters the vascular endothelial cell redox state and its potency relative to known mitochondrial antioxidants. Another noteworthy observation of this study was that EPI did not rescue AA-induced increases in mitochondrial ROS. Likewise, one recent study reported that 1 µM EPI did not lower the production of mitochondrial superoxide in HUVECs after acute AA treatment (Keller et al., 2020), suggesting limited efficacy for EPI in overcoming conditions associated with elevated mitochondrial ROS production in the vascular endothelium of humans.

The potent stimulation of NO production by EPI is well documented in studies using cell, human and rodent models (Loke et al., 2008; Moreno-Ulloa, Mendez-Luna, et al., 2015; Ramirez-Sanchez et al., 2011, 2018; Schroeter et al., 2006), although not all studies have reported such effects (Dower et al., 2015). Here, we demonstrated that EPI increased NO levels of vascular endothelial cells, suggesting that EPI may be a promising strategy to combat vascular endothelial dysfunction, although further *in vivo* studies are required. Previous studies have attributed elevated NO bioavailability in the presence of EPI to increased phosphorylation of eNOS at Ser177 (Carnevale et al., 2014; Ramirez-Sanchez et al., 2010, 2012, 2018; RamÍrez-Sánchez et al., 2016). Given that we and others demonstrated unaltered eNOS phosphorylation in the presence of EPI (Keller et al., 2020), it is plausible that arginase inhibition is the potential mechanism by which EPI increases NO production in HUVECs (Schnorr et al., 2008). However, this may not be the case in arterial endothelial cells, where EPI has repeatedly been shown to enhance eNOS Ser1177 phosphorylation (Ramirez-Sanchez et al., 2012, 2018; RamÍrez-Sánchez et al., 2016).

### EPI augments the expression of genes linked with the antioxidant response and mitochondrial dynamics

To better understand the mechanisms underlying EPI’s effects, we measured the transcription of genes associated with energy metabolism. Interestingly, EPI increased the expression of genes DRP1 and MFN2, respectively involved in mitochondrial fission and fusion. Despite these effects, EPI did not impact mitochondrial respiration, at least over 24 hours. Although mitochondrial dynamics can influence respiratory function (Chen et al., 2010; Glancy et al., 2020), the lack of functional change in mitochondrial respiration could reflect a discord between cellular mRNA responses and changes in protein abundance and/or function. Regardless of dose, EPI significantly enhanced NRF2 mRNA expression. This observation implies that the induction of NRF2 mRNA by EPI is independent of RONS production and might be explained by alternate factors that control NRF2 activity, like phosphorylation status (Robledinos-Antón et al., 2019). Although post-translational modifications of NRF2 were not assessed in this study, flavonoids have been shown to promote NRF2 phosphorylation and nuclear translocation (Lan et al., 2017; Shi et al., 2018), which would likely increase the transcription of genes related to the antioxidant response, including NRF2 itself.

### EPI does not impact mitochondrial respiration

One important finding was that EPI does not impact mitochondrial respiration of HUVECs in culture. Although mitochondria have been proposed as potential molecular targets of EPI (Duluc et al., 2012), previous investigations into EPI’s effects on mitochondrial respiration have produced equivocal results. Some studies have demonstrated increased state 3 respiration in rat beta cells following EPI (0.1-2.5 µM) supplementation (Kener et al., 2018; Rowley et al., 2017), whilst others have reported inhibited or similar state 3 respiration rates with EPI, depending on the substrates provided (Kopustinskiene et al., 2015). One recent study using HUVECs as a model reported that 0.1 and 1 µM EPI treatment over 2 hours had negligible impact on mitochondrial respiration (Keller et al., 2020). Taken together, our data suggest that the therapeutic potential of EPI is not related to changes in mitochondrial respiration in vascular endothelial cells, pointing to alternate potential mechanisms of action like cell signalling activation (Fraga et al., 2018).

### EPI rapidly and transiently activates ERK1/2 signalling

The serine/threonine protein kinase AMPK is an important regulator of mitochondrial adaptation (Herzig & Shaw, 2018). However, in our studies AMPK phosphorylation was suppressed in the presence of EPI for up to 1 h and was without further impact for up to 24 h, compared with controls. Supporting these findings, 2 h EPI treatment (1 µM) did not affect the phosphorylation of AMPK in HUVECs (Keller et al., 2020). Although EPI is capable of augmenting AMPK activity in liver and muscle tissue, and several cell types (Murase et al., 2009; Papadimitriou et al., 2014; Si et al., 2011), it seems that AMPK activation is not responsible for the therapeutic actions of EPI in HUVECs in culture. Together with the negligible impact of EPI on mitochondrial respiration, the data suggest that EPI does not directly alter vascular endothelial cell energy metabolism, *in vitro*.

Importantly and of novelty, we reported that ERK1/2 phosphorylation at Thr202/Tyr204 was rapidly and transiently increased by EPI in HUVECs (Figure 5). This finding supports recent observations of increased ERK1/2 signalling after 0.1 µM EPI treatment in bovine coronary artery endothelial cells, that may be associated with phosphorylation of CaMKII (Moreno-Ulloa, Mendez-Luna, et al., 2015). How EPI promotes ERK1/2 phosphorylation in vascular endothelial cells remains to be described, but current evidence suggests EPI activates ERK1/2 by binding to the G-protein coupled estrogen receptor (GPER) on the cell membrane (Moreno-Ulloa, Mendez-Luna, et al., 2015; Yang & Chan, 2018). Given the induction of NRF2 mRNA expression found in this study, it would be useful to determine if acute activation of ERK1/2 signalling by EPI is a prerequisite for the induction of NRF2 activity.

## Limitations

We did not use metabolites of EPI that appear in circulation after ingestion of EPI-containing foods or supplements *in vivo* (Ottaviani et al., 2016). Therefore, the present findings may have limited translational potential. In our study we harnessed HUVECs to model the vascular endothelial cell, and given that these cells are venous in nature, their physiology may not well reflect the arterial vasculature or microcirculation where EPI potentially exerts its beneficial effects. Further, not all assays were performed in the presence of additional cell stress. Thus, caution should be taken when translating our findings to human populations with disease. Finally, post-translational modifications of NRF2, which are critical for regulating NRF2’s activity, were not measured in this study.

## Conclusion

We demonstrate that physiologic EPI concentrations do not impact mitochondrial respiration but do modulate reactive oxygen and nitrogen species production and the signalling and transcriptional activities of vascular endothelial cells *in vitro*. EPI’s dose-dependent alteration of reactive oxygen and nitrogen species production occurred in parallel with enhanced and transient ERK1/2 signalling, and the induction of NRF2 mRNA (Figure 6). The fact that EPI enhanced NRF2 mRNA expression regardless of dose, implies that alterations in reactive oxygen and nitrogen species production alone were not solely responsible. Further research will help clarify the precise way in which EPI promotes ERK1/2 signalling and NRF2 activity, and its relevance to vascular endothelial health *in vivo*.

**Figure 6.**
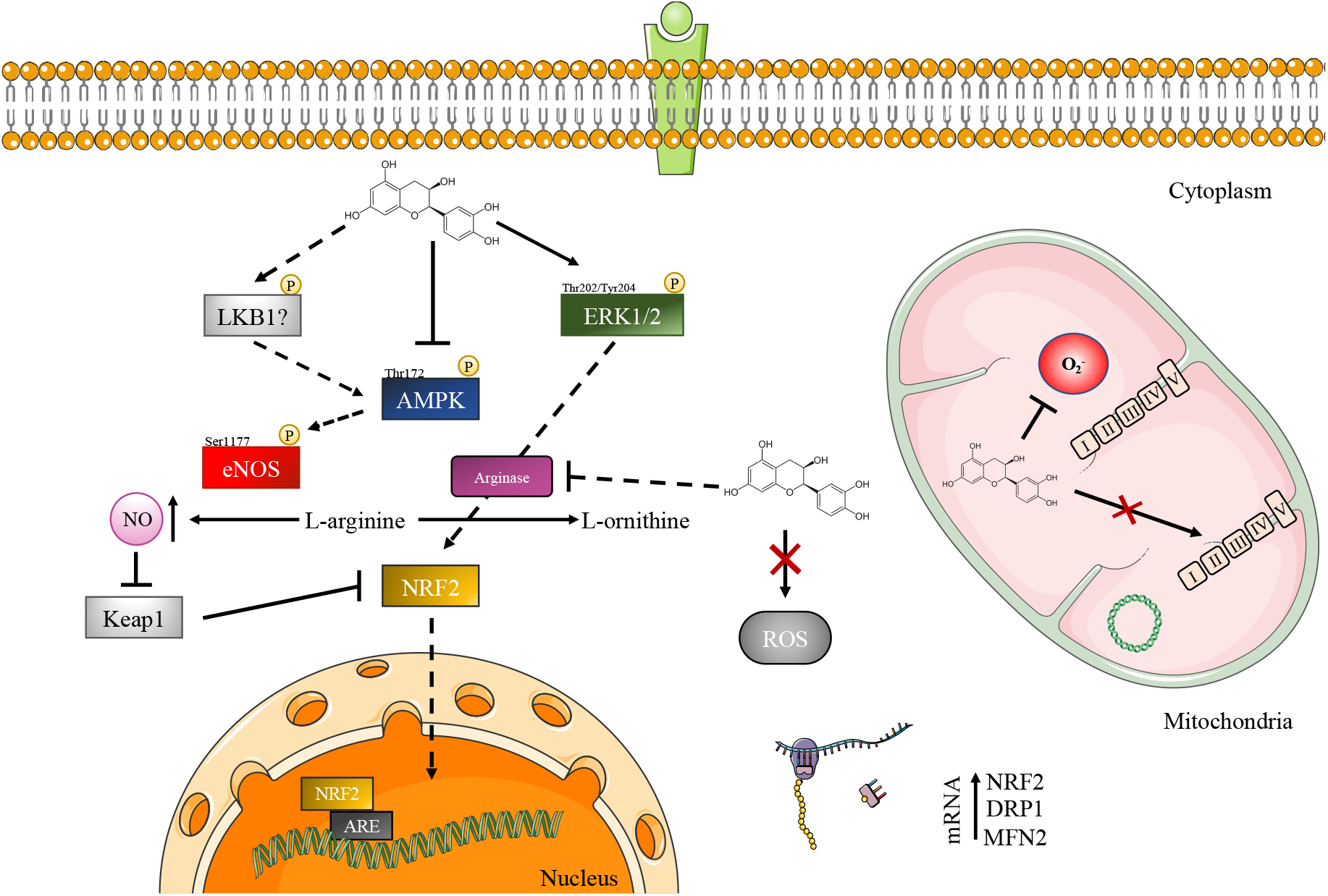
Schematic of the potential mechanisms by which EPI exerts its biological effects in vascular endothelial cells. Solid arrows/lines signify EPI’s mode of action demonstrated in this study. Dashed arrows represent potential activity of EPI not examined in this study but reported previously.

## Availability of data and material

The datasets generated and analysed during this study are available from the corresponding author upon request.

## Authors’ contributions

DGS and CES conceived the study and designed experiments. DGS and JB designed the respiration experiments using the Seahorse XFe96 Analyzer. DGS collected and analysed the data, DGS, JB, RD, HJ, DHJT and CES interpreted the data. DGS and CES wrote the manuscript and all authors revised it critically. All authors provided final approval of the version to be published and agree to be accountable for all aspects of the work in ensuring that questions related to the accuracy or integrity of any part of the work are appropriately investigated and resolved. All people designated as authors qualify for authorship, and all those who qualify for authorship are listed. CES is the guarantor for the work and/or conduct of the study, had full access to all the data in the study and takes responsibility for the integrity of data and the accuracy of the data analysis, and controlled the decision to publish.

## Acknowledgements

The authors acknowledge use of the Mitochondrial Profiling Centre, a core resource supported by the University of Birmingham. This study was supported by the BBSRC funded Industrial CASE (iCASE) award (BB/P504385/1).

