## Supplementary Information for "(–)-Epicatechin alters reactive oxygen and nitrogen species production independent of mitochondrial respiration in human vascular endothelial cells"

**Table S1.** Primer sequences for homo sapiens with product length. All primers were used under the same cycling conditions.

| Gene | Accession | Sequence<br>Forward/Reverse or Anchor Nucleotide | Product length (bp) |
|---|---|---|---|
| RPL13a | NM_012423.4 | F: GGCTAAACAGGTACTGCTGGG<br>R: GGAAAGCCAGGTACTTCAACT | 104 |
| CAT | NM_001752 | AN: 1649 | 134 |
| SOD2 | NM_000636 | AN: 194 | 132 |
| DNM1L (DRP1) | NM_012062.5 | F: CACCCGGAGACCTCTCATTC<br>R: CCCCATTCTTCTGCTTCCAC | 99 |
| MFN2 | NM_014874.4 | F: CCCCCTTGTCTTTATGCTGATGT<br>R: TTTTGGGAGAGGTGTTGCTTATT | 168 |
| PPARGC1A (PGC-1α) | NM_001330751.2 | F: TGCTAAACGACTCCGAGAA<br>R: TGCAAAGTTCCCTCTCTGCT | 67 |
| SIRT1 | NM_012238 | AN: 1382 | 109 |
| TFAM | NM_003201 | AN: 462 | 143 |
| NOS3 (eNOS) | NM_000603.5 | F: AACTATTTCCTGTCCCCGGC<br>R: AGGATTGTCGCCTTCACTCG | 173 |
| CYBB (NOX2) | NM_000397.4 | F: GGGCTGTTCAATGCTTGTGG<br>R: GGCCCATCAACCGCTATCTT | 80 |
| NOX4 | NM_016931.5 | F: CAGTCCTTCCGTTGGTTTGC<br>R: CAAAAGTTTCCACCGAGGACG | 189 |
| PRKN (PARKIN) | NM_004562 | AN: 747 | 91 |
| GABPA (NRF2) | NM_002040.4 | F: AAATTGAGATTGATGGAACAGAGAA<br>R: TATGGCCTGGCTTACACATTCA | 95 |

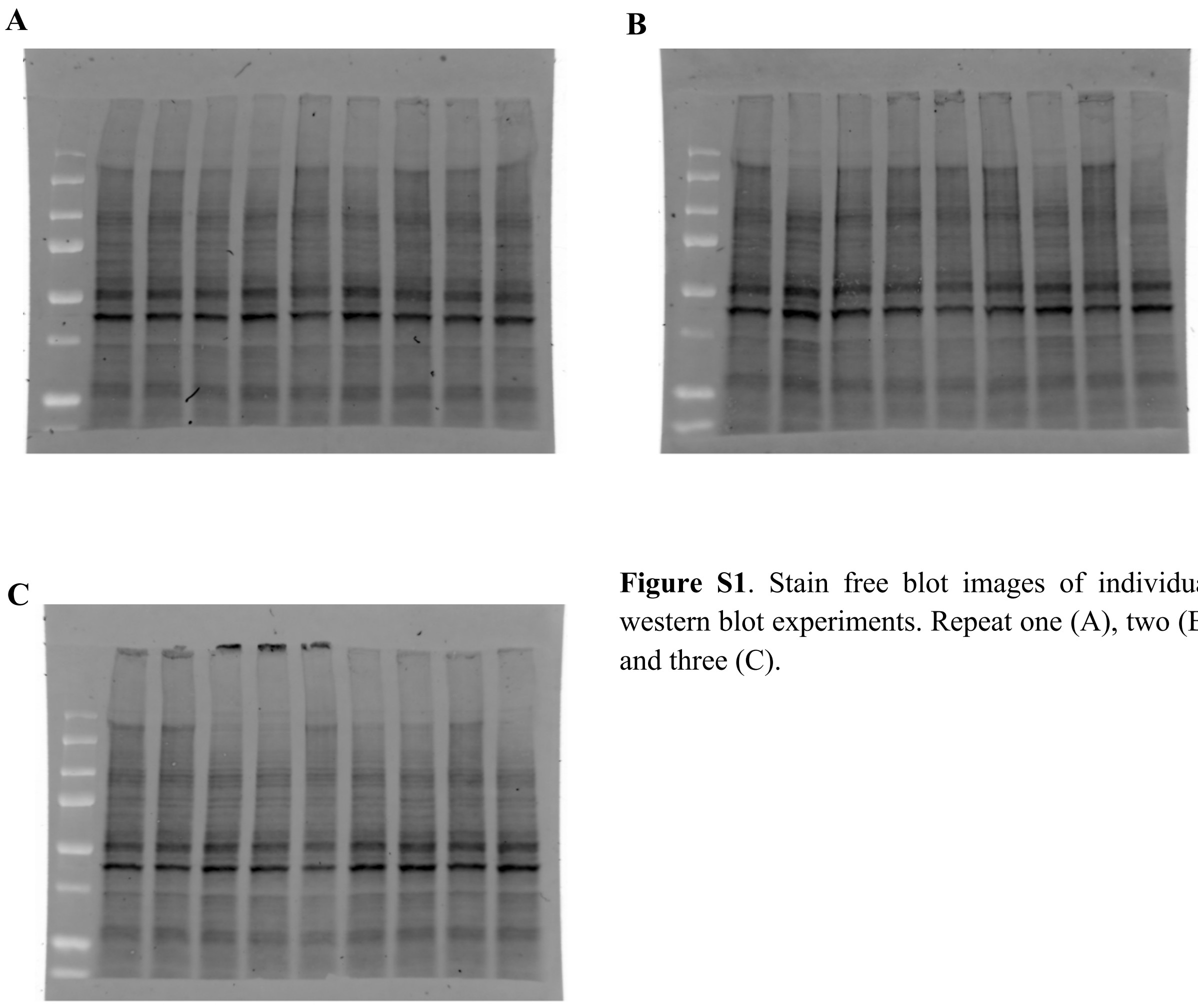


**Figure S1**. Stain free blot images of individual western blot experiments. Repeat one (A), two (B) and three (C).

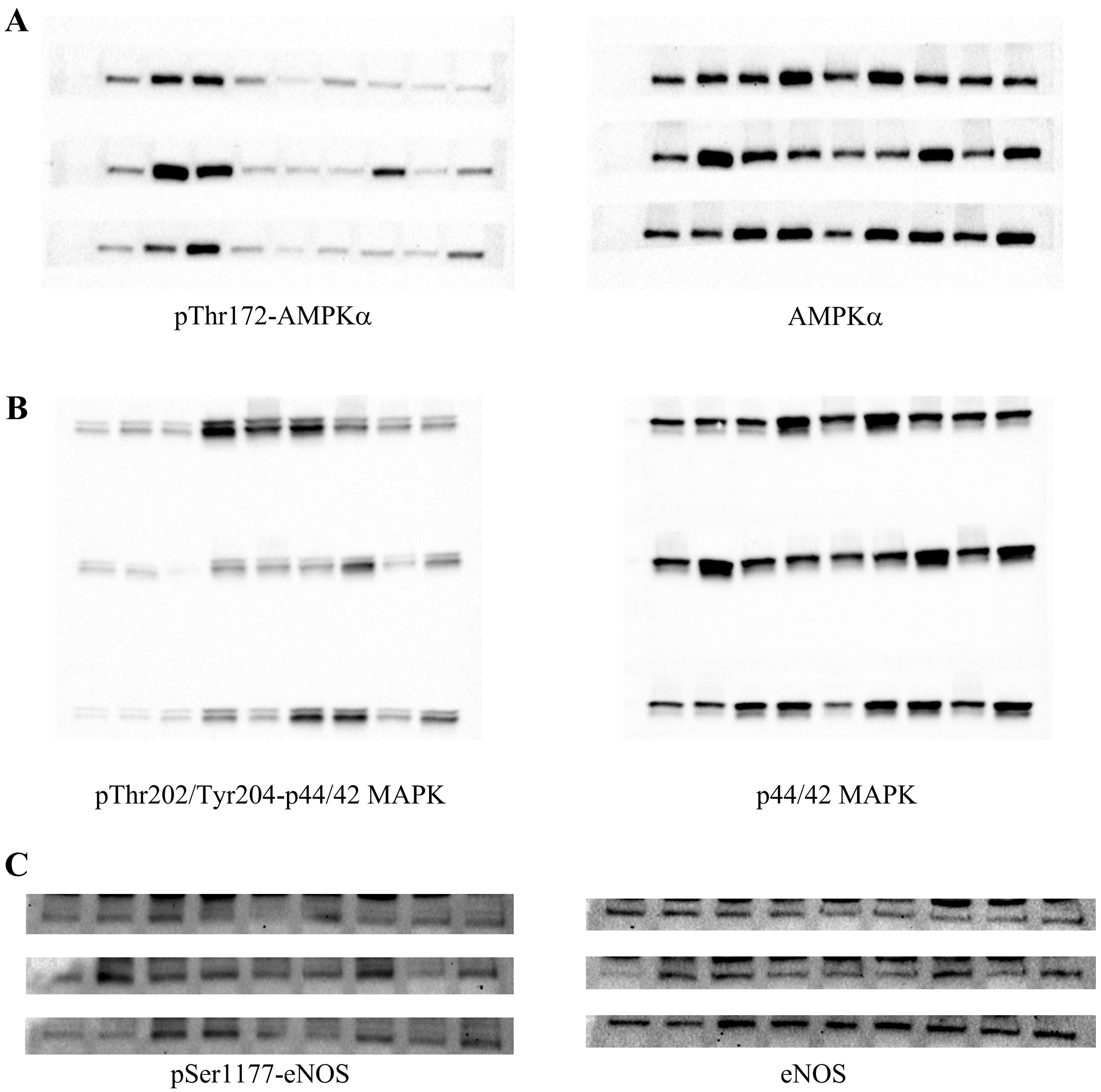


**Figure S2**. Western blot analysis of HUVEC lysates. A) pThr172-AMPKα and total AMPKα, B) pThr202/Tyr204-p44/42 MAPK and total p44/42 MAPK, C) pSer1177-eNOS and total eNOS.

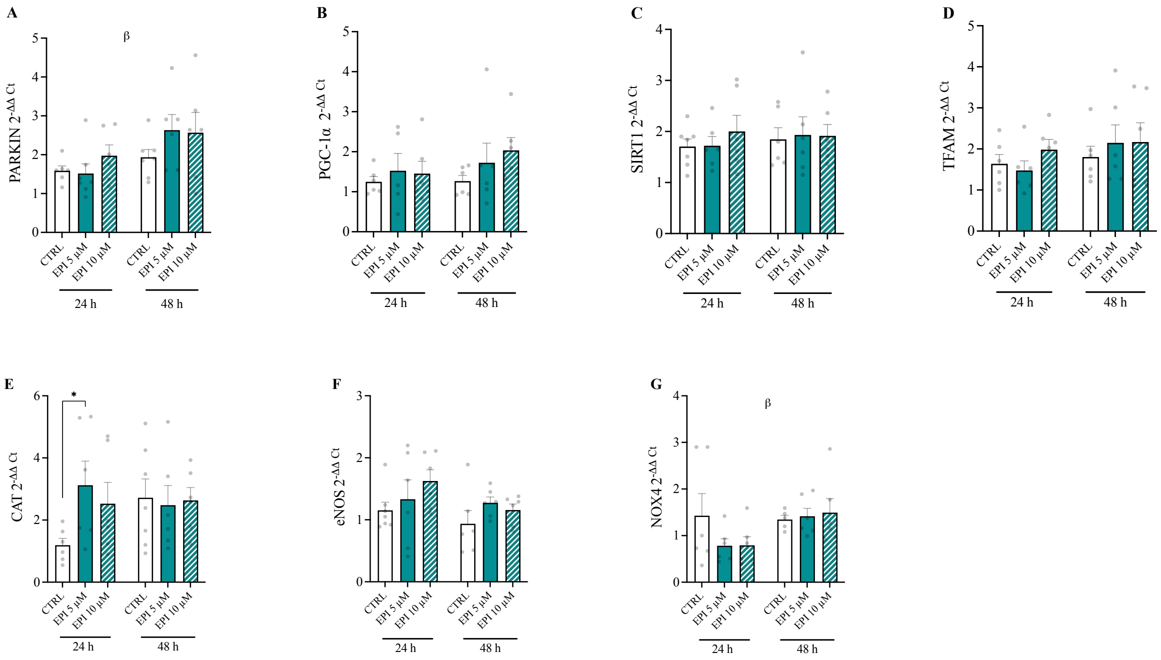


**Figure S3**. Gene expression responses following acute EPI treatment. HUVECs were treated with 0, 5 and 10 μM EPI over 48 h and lysed for analysis of gene expression. A) Parkin, B) PGC-1α, C) Sirt1, D) Tfam, E) Catalase, F) eNOS and G) NOX4. Data are means±SEM from 3 independent experiments. Statistical significance was determined by a two-way ANOVA, with dose and time as factors. Multiple comparisons were performed by Dunnett's test to determine differences in gene expression between conditions. $^{\beta}$ main effect of time ($P<0.05$); *$P<0.05$.
